# CRISPR-based tools for genetic manipulation in pathogenic *Sporothrix* species

**DOI:** 10.1101/2022.11.15.516583

**Authors:** Remi Hatinguais, Ian Leaves, Gordon D. Brown, Alistair J. P. Brown, Matthias Brock, Roberta Peres da Silva

## Abstract

*Sporothrix brasiliensis* is an emerging fungal pathogen frequently associated with zoonotic transmission of sporotrichosis. Although certain virulence factors have been proposed as potential sporotrichosis determinants, the scarcity of molecular tools for reverse genetics studies on *Sporothrix* has significantly impeded the dissection of mechanisms underlying the disease. Here, we demonstrate that PEG-mediated protoplast transformation is a powerful method for heterologous expression in *S. brasiliensis, S. schenckii* and *S. chilensis*. Combined with CRISPR/Cas9 gene editing, this transformation protocol allowed the deletion of the putative DHN-melanin synthase gene *pks1*, which is a proposed virulence factor of *Sporothrix* species. To improve in locus integration of deletion constructs, we deleted the KU^80^ homologue that is critical for non-homologous end-joining DNA repair. The use of *S. brasiliensis* Δ*ku80* strains enhanced homologous-directed repair during transformation resulting in increased targeted gene deletion. In conclusion, our CRISPR/Cas9-based transformation protocol provides an efficient tool for targeted gene manipulation in *Sporothrix* species.

## INTRODUCTION

Sporotrichosis is a neglected fungal disease caused by thermo-dimorphic fungi belonging to the *Sporothrix* complex, which includes *S. brasiliensis, S. schenckii* sensu stricto, *S. luriei* and *S. globosa*. While those species are frequently associated with human and animal disease, the environmental species *S. mexicana, S. pallida*, and *S. chilensis* are referred to as opportunistic pathogens that cause infections in immunocompromised patients (Orofino-Costa et al., 2017; Rodrigues et al., 2022). Sporotrichosis is classically transmitted by traumatic inoculation of fungal propagules into subcutaneous tissue (Orofino-Costa et al., 2022). Nevertheless, ongoing outbreaks in Brazil are mainly related to the epizootic transmission among cats infected with *S. brasiliensis*, a species of high virulence. Due to their close proximity to humans, zoonotic transmission from cats via scratches or bites has become a challenge for confining the geographical advance of the disease (Garcia Carnero et al., 2018; Orofino-Costa *et al*., 2022; Orofino-Costa *et al*., 2017). Over about 25 years, a sporotrichosis outbreak that started in the state of Rio de Janeiro has spread throughout almost all the Brazilian territory and into few of its neighbouring countries in Latin America, such as Argentina (Etchecopaz et al., 2021; Orofino-Costa *et al*., 2022). Furthermore, at least one case of sporotrichosis, transmitted by an imported infected cat, has been reported in the United Kingdom (Rachman et al., 2022). Thermo-dimorphism, glycans, adhesins, secreted vesicles and melanin have all been associated with the ability of the fungus to cause disease in the mammalian host (Garcia-Carnero and Martinez-Alvarez, 2022; Garcia Carnero *et al*., 2018). However, their contributions to virulence remain to be confirmed experimentally.

*S. brasiliensis* and *S. schenckii* possess three pathways related to melanin production: 3,4-dihydroxy-L-phenylalanine (l-DOPA), 1,8-dihydroxynaphthalene (DHN)-melanin and pyomelanin (Almeida-Paes et al., 2017; Teixeira et al., 2014). The presence of melanin in fungal cells has been related to resistance to environmental stresses and the survival of the pathogen during host interactions (Almeida-Paes *et al*., 2017; Cordero and Casadevall, 2017; Liu et al., 2021). *S. schenckii* yeasts and conidia produce melanin particles that can be recognized by monoclonal antibodies produced from murine hybridomas and sera from patients with sporotrichosis (Morris-Jones et al., 2003). UV-induced DHN-melanin deficient strains of *S. schenckii* were less resistant to oxidative/nitrosative stresses and phagocytosis by human monocytes or murine macrophages (Romero-Martinez et al., 2000). Conidia from a wild-type (WT) *Sporothrix globosa* strain revealed a lower expression of MHC class II in murine macrophages and a higher fungal burden in spleen when compared to an UV-induced albino mutant (Guan et al., 2021; Masternak et al., 2000; Song et al., 2021). In addition, DHN-melanin inhibitors enhanced the *in vitro* susceptibility of *S. brasiliensis* and *S. schenckii* to the antifungal drug terbinafine. In addition, yeast cells from *S. brasiliensis* and *S. schenckii* grown in the presence of the melanin-inducing L-DOPA or L-tyrosine were less susceptible to amphotericin B, but not itraconazole (Almeida-Paes et al., 2016; Mario et al., 2016). In contrast, the inhibition of the eumelanin or pyomelanin pathways did not result in an increased antifungal susceptibility (Almeida-Paes *et al*., 2016). This indicates the general importance of all three types of melanin during host infection and the particular importance of DHN-melanin in promoting resistance to antifungal therapy.

In the last two decades, there has been an increased effort to develop new tools for diagnosis, epidemiologic and virulence studies, but few studies have focused on new approaches to improve the molecular and genetic manipulation of the *Sporothrix* species complex (de Carvalho et al., 2022; Rodrigues *et al*., 2022). Currently, the genetic manipulation of *Sporothrix* species has focused on gene silencing using vectors delivered to fungal cells *via Agrobacterium tumefaciens-mediated* transformation, or by random UV-induced mutation of *S. schenckii* and *S. globosa* (Lozoya-Perez et al., 2019; Lozoya-Perez et al., 2018; Romero-Martinez *et al*., 2000; Song et al., 2021; Tamez-Castrellon et al., 2018; Tamez-Castrellon et al., 2021). *Sporothrix* species have represented a challenge for the application of universal genetic engineering tools (Mora-Montes et al., 2015). This has hindered the elaboration of specific mechanisms that promote the survival and immune evasion of *Sporothrix* species during host interactions.

In this study, we have established a protocol for accurate gene disruption in *S. brasiliensis* and *S. schenckii* using the CRISPR/Cas9 system. Furthermore, we have standardised a transformation protocol for ectopic integration and heterologous gene expression in *S. chilensis*, a species without a sequenced genome. Moreover, we show that the deletion of the *ku80* gene, together with the functional expression of a red-shifted luciferase in *S. brasiliensis* strains, facilitate the screening of transformants and provide a promising tool for *in vivo* real-time monitoring of sporotrichosis.

## RESULTS

### Protoplast-mediated transformation grants delivery and ectopic integration of synthetic genes in pathogenic *Sporothrix* species

Protoplast mediated transformation of filamentous fungi is a well-established method (Li et al., 2017), but such an approach has not been explored for its suitability on the mycelium phase of thermo-dimorphic fungi, such as *Sporothrix* species. To develop an efficient protocol for transformation in *Sporothrix* species, our first step was to assess appropriate combinations of growth medium and selection markers that could be utilised during PEG-mediated protoplast transformation. We performed spot assays of serial dilutions of spores from the strains *S. brasiliensis* Ss54*, S. schenckii* Ss126 and *S. chilensis* Ss469. These strains were tested for the ability to grow in modified *Aspergillus* minimal medium containing 50 mM glucose as the carbon source, 10 mM glutamine as the nitrogen source (GG10) (Geib and Brock, 2017) and supplemented with thiamine (GG10_THI_). Sorbitol was added as an osmotic stabilizer of protoplasts. In addition, we tested three different concentrations of antibiotics for two selection markers previously utilized in *S. schenckii* transformation (Lozoya-Perez et al., 2019; Tamez-Castrellon et al., 2018): hygromycin B (80, 140 and 180 μg/ml) and nourseothricin (50, 100 and 200 μg/ml). The concentrations of 140 μg/ml hygromycin B and 50 μg/ml nourseothricin were sufficient to inhibit mycelial growth of all strains tested (Figure 1A).

**Figure 1:**
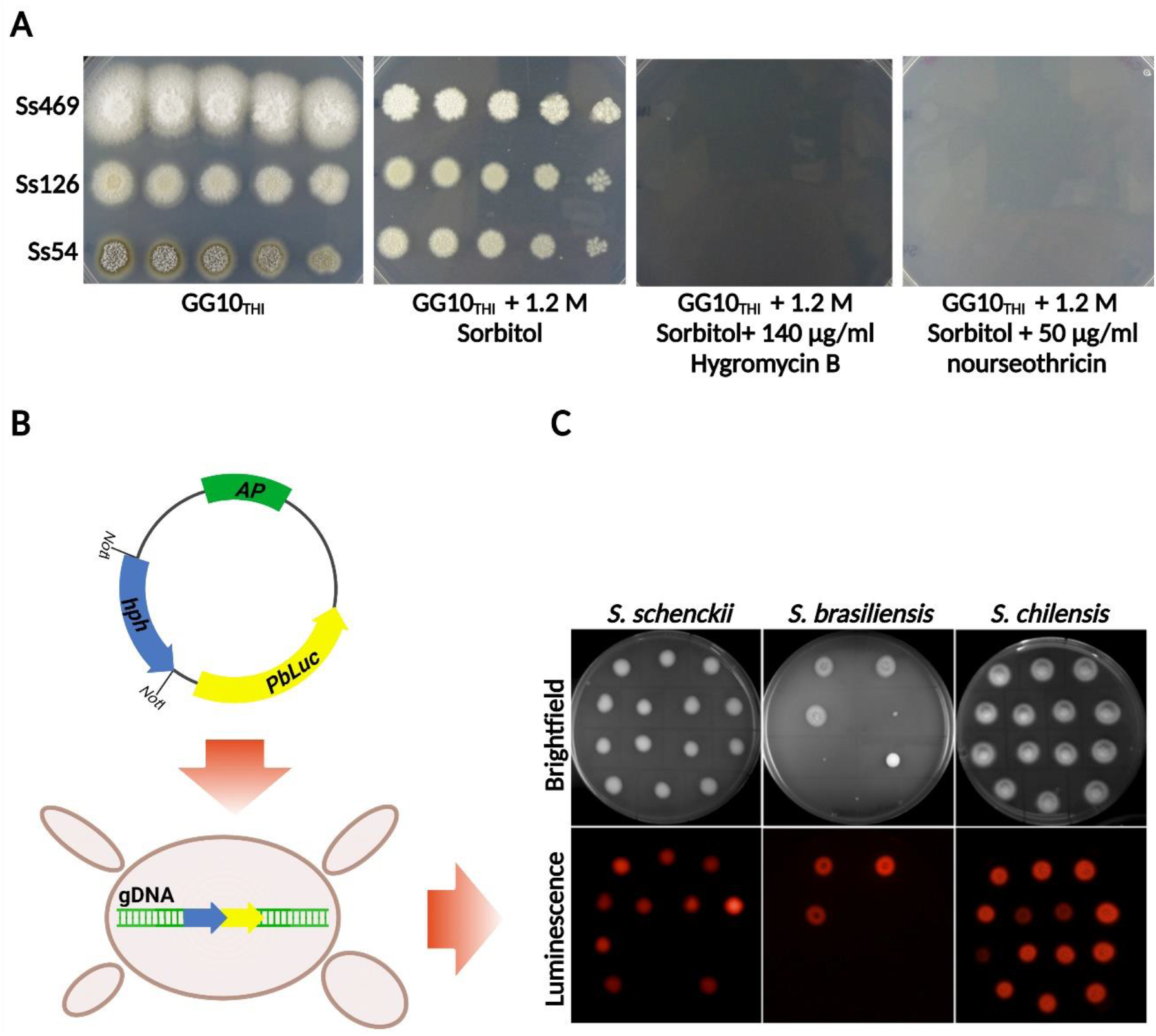
Protoplast-mediated transformation of *S. schenckii, S. brasiliensis* and *S. chilensis*. **(A)** Spot assays of 5 serial dilutions ranging from 3 × 10^5^ to 30 spores from *S. brasiliensis* Ss54*, S. schenckii* Ss126 and *S. chilensis* Ss469 on GG10_THI_, GG10_THI_ supplemented with 1.2 M sorbitol, 140 μg/ml hygromycin B and 50 μg/ml nourseothricin. **(B)** Schematic representation of the plasmid employed in the transformation protocol showing the *hph* resistance cassette and red-shifted firefly luciferase (*PbLuc*) construct. **(C)** Imaging of transformants containing the luciferase construct inoculated onto agar plates containing Hygromycin B (140 μg/ml) and d-luciferin (0.2 mM). Brightfield (upper panels) and luminescence (bottom panels) were recorded after 7 days of incubation.

To test for the stable genomic integration of DNA constructs, we utilized a plasmid harbouring the synthetic coding region for a thermostable red-shifted luciferase that was previously codon-optimized for use in *Paracoccidioides brasiliensis* (*PbLuc*) (Milhomem Cruz-Leite et al., 2022) and expressed under the control of *Paracoccidioides* promoters from the *elongation factor 1-gamma* (*ef1*), *enolase 1* (*eno1*) or *actin* (*act*) genes. These plasmids had been shown to drive functional luciferase expression in both *Aspergillus niger* and *Paracoccidioides* species (Supplementary Figure 1A-D and Milhomem Cruz-Leite *et al*. (2022)).

Protoplasts from *Sporothrix* mycelia were generated by enzymatic digestion of cell wall components using logarithmically growing cells (Li *et al*., 2017). The fusion of protoplasts and the uptake of the plasmid harbouring *Pef1:PbLuc_hph, Peno1:PbLuc_hph or Pact:PbLuc_hph* was mediated by polyethylene glycol (PEG) transformation as previously described for aspergilli (Figure 1B) (Geib and Brock, 2017; Gressler et al., 2015; Peres da Silva and Brock, 2022). This procedure resulted in several bioluminescent transformants for all three strains tested (Ss54, Ss126 and Ss469). It is worth noting that, using this protocol, *S. brasiliensis* transformations always resulted in lower numbers of transformants compared to the other strains tested (Figure 1C). A selection of bioluminescent transformants was passaged at least 10 times in the absence of selective pressure. The insertion of luciferase into the genome and its expression proved to be stable for at least three years thereby confirming the genetic stability of transformants.

### CRISPR/Cas9 gene editing is an efficient approach for gene disruption in *Sporothrix*

*S. schenckii* and *S. brasiliensis* produce colonies with brownish mycelia when cultivated on solid medium, due the accumulation of melanin pigments formed *via* the DHN-melanin pathway (Almeida-Paes *et al*., 2016; Almeida-Paes et al., 2009; Romero-Martinez *et al*., 2000). We identified the putative polyketide synthase gene (*pks1;* SPSK_00653 and SPBR_06313) likely to be responsible for the production of 1,3,6,8-tetrahydroxynaphthalene (THN) on this DHN-melanin pathway in *Sporothrix*, due to its sequence similarity to the *Colletotrichum lagenarium pks1* gene (Fujii et al., 1999) and the *A. fumigatus* naphthopyrone synthase *pksP* gene (Langfelder et al., 1998; Watanabe et al., 2000). We then targeted *pks1* for gene deletion using the CRISPR/Cas9 system.

Deletion of the *pks1* gene was predicted to eliminate the biosynthesis of DHN-melanin resulting in white mycelia. This loss of mycelial colour was used to pre-select for transformants in which the *pks1* gene had been successfully inactivated. Regions of about 850 bp of flanking homology lying upstream and downstream of the Cas9 enzyme cleavage sites in the *Sporothrix pks1* gene were amplified and fused with nourseothricin resistance cassette (Figure 2A and Supplementary table 1). Guide RNAs (gRNAs) complexed with Cas9 were used to create the *pks1* gene deletion in Ss126 *S. schenckii*, MYA-4823 and Ss54 *S. brasiliensis* wild-type (WT) strains. Two different gRNAs were designed to cut at about 29 bp and 2199 bp after the predicted ATG start codon, corresponding to an excision of the first protein domain plus part of the second domain of the putative polyketide synthase Pks1. The excised *pks1* fragment was replaced by a stop codon sequence plus a nourseothricin resistance cassette of approximately 1.7 kb excluding the homology arms.

**Figure 2:**
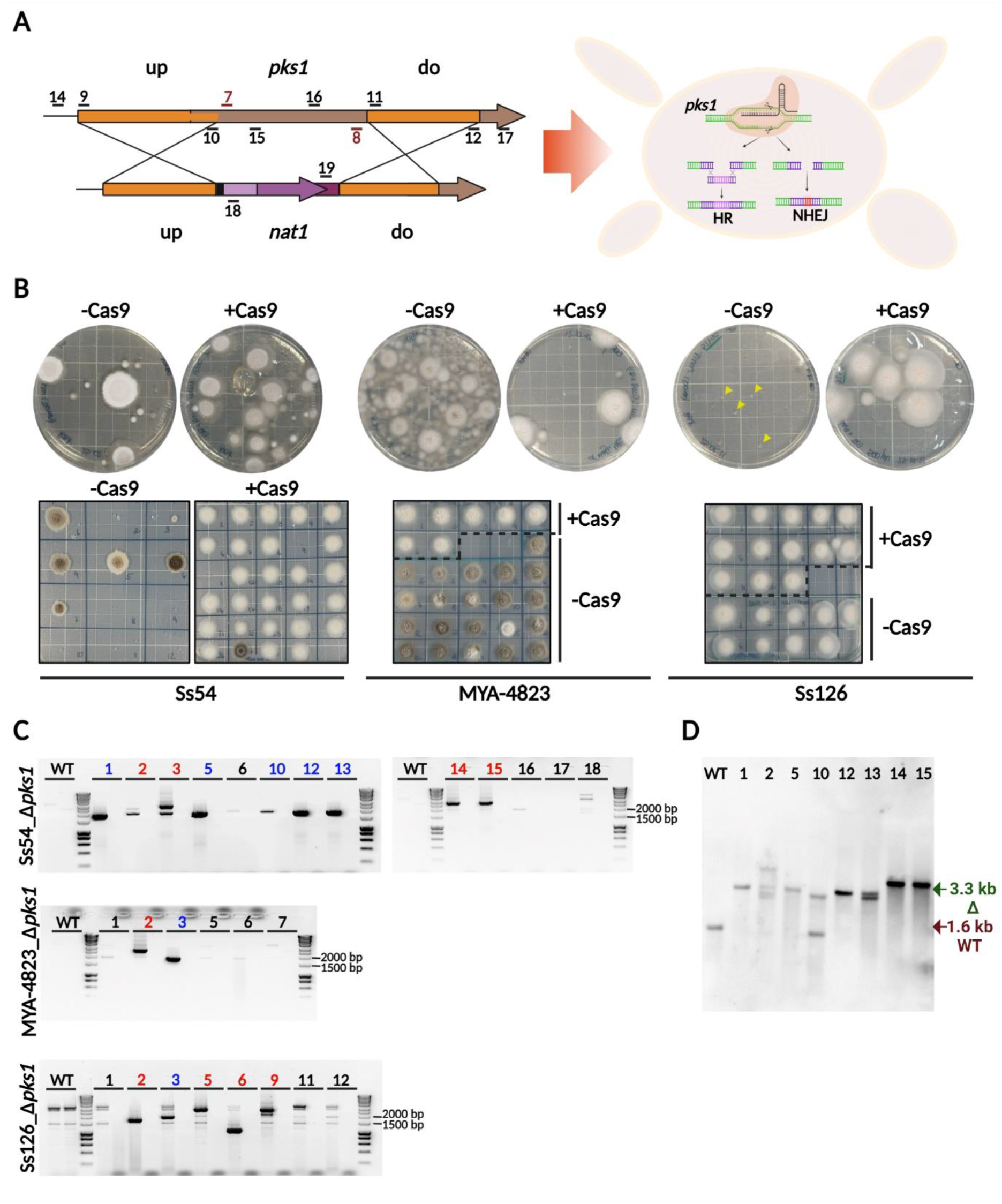
Disruption of the *pks1* gene. **(A)** Schematic representation of the *pks1* deletion construct indicating: the annealing position of oligonucleotides (black) and gRNAs (red) utilized during the gene disruption and transformants selection; the crossover regions predicted for homologous recombination; the homologous recombination (HR) and non-homologous end-joining (NHEJ) pathways for repair of DNA double-strand breaks active during gene disruption. **(B)** Representative transformation agar plates for *Sporothrix* strains Ss54, MYA-4823 and Ss126 (above) along with subculture of putative *pks1* transformants onto GG10_THI_ containing nourseothricin 80 μg/ml (below). **(C)** PCR analysis for detection of the partial replacement of *pks1* coding region by the deletion cassette containing nourseothricin resistance gene. The insertion of the deletion cassette via homology-directed repair in the upstream flanking region resulted in PCR products of 2000 bp (samples highlighted in blue), or insertion via non-homologous repair mechanisms (PCR products ≠ 2000 bp, samples highlighted in red). **(D)** Southern blot analysis of Ss54 transformants that showed integration of the deletion cassette on the upstream flanking region during the PCR screening. Diagnostic bands of 1.6 kb for wild type (WT) and 3.3 kb for a deletion mutant (Δ) are shown.

Transformations were performed with or without Cas9 in parallel to evaluate whether the flanking homologies from the donor DNA are sufficient for targeted gene disruption. During transformation of *S. schenckii* Ss126, the absence of Cas9 resulted in colonies with a smaller size on the transformation agar. However, these transformants showed the expected growth pattern in GG10_THI_ supplemented with nourseothricin (Figure 2B), indicating that these transformants had incorporated the deletion cassette into the genome.

Transformations using Cas9 were more efficient in terms of the yields of putative transformants for *S. brasiliensis* Ss54, but not for *S. brasiliensis* MYA-4823. Interestingly, none of the strains generated brown colonies on the transformation agar, suggesting that the *pks1* gene had been successfully inactivated in the presence or absence of Cas9. Nonetheless, the transfer of colonies to GG10_THI_ medium lacking sorbitol and supplemented with nourseothricin did result in the development of a brownish colour for some Ss54 and MYA-4823 transformants, mostly for colonies generated during the transformation without Cas9 (Figure 2B). Interestingly, none of colonies from the *S. schenckii* Ss126 transformation developed brownish colour in GG10_THI_ or potato dextrose agar (PDA), suggesting successful deletion of *pks1*.

To evaluate the efficiency of the gene disruption, we performed diagnostic PCR analyses on genomic DNA (gDNA). Oligonucleotides flanking the upstream and downstream regions of the excised *pks1* sequence were used to differentiate between the genotypes of WT strains and *pks1* deleted transformants. We tested 16 Ss54 transformants, 7 MYA-4823 transformants and 12 Ss126 transformants that were generated using Cas9 (Supplemental Figure 2). We also tested 5 Ss54 colonies, 5 Ss126 colonies and 12 MYA-4823 colonies that were derived from transformations lacking Cas9 and receiving only donor DNA (Supplemental Figure 2). The WT coding region yielded PCR products of 2257 bp and 2337 bp for the upstream and downstream regions, respectively, whereas *pks1* deletion or recombination were assumed to produce no PCR products. About 75% of the transformants generated using Cas9 showed the expected disruption of the *pks1* coding sequence (samples highlighted in purple in Supplementary Figure 2). On the other hand, the *pks1* gene was only disrupted in its downstream region in the majority of Ss54 and MYA-4823 transformants generated without Cas9 (samples highlighted in green in Supplementary Figure 2). This suggested that instead of recombination events taking place at both, the upstream and downstream flanking homologous regions used to replace the *pks1* coding sequence with the disruption cassette only a single recombination event had taken place in the absence of Cas9. This resulted in a predominant insertion of the deletion cassette at the downstream flanking homology arm. This type of event did not result in white colonies (Figure 2B and Supplemental Figure 2).

Those colonies from the Cas9-containing transformations in which the *pks1* coding sequence was no longer detected in both PCRs were tested further by positional PCR for the integration of the nourseothricin cassette within the *pks1* locus (Figure 2C). The insertion of the disruption cassette within the *pks1* locus mediated by homology-directed repair resulted in PCR products of 2000 bp and 1647 bp for the upstream and downstream regions, respectively, while an intact WT *pks1* coding region was not expected to result in PCR products. The nourseothricin cassette was detected in nine of the thirteen Ss54 transformants tested. However, in all positive samples the recombination only occured in the upstream region: seven via precise homologous recombination (to yield the 2000 bp PCR product; samples highlighted in blue), and two via non-homologous DNA repair mechanisms (to yield a PCR product that was not 2000 bp; samples highlighted in red). For the MYA-4823 strain, one transformant had the donor DNA inserted into the genome by homologous recombination, one had integrated the DNA by non-homologous processes, and no integration of the nourseothricin cassette was detected in the five remaining transformants, indicating an ectopic integration of the *pks1* deletion cassette that was not further analysed in this PCR screening. For the eight transformants obtained for Ss126, only one had the nourseothricin cassette integrated at the *pks1* locus by homologous recombination.

Southern blot analysis was performed on Ss54 transformants to test whether the donor DNA had integrated only at the target locus, or ectopically at additional sites (Figure 2D). The majority of the transformants tested showed a single integration event. The data indicated that CRISPR/Cas9 promotes efficient targeted mutation in *Sporothrix*. However, the donor DNA integration events seemed to appear at a preferred individual Cas9 site, indicating that gene disruption was more likely to occur compared to a gene deletion event. As expected, the disruption of the *pks1* locus resulted in white colonies, showing that the gene is required for the brown colouration of mycelia. Unfortunately, a significant proportion of the transformants resulted from non-homologous integration events, which reduced the number of transformants carrying a true *pks1* gene deletion in *Sporothrix* species.

### Enhancing homologous recombination in *S. brasiliensis*

Our results showed that the *pks1* gene from *Sporothrix* can be successfully disrupted by applying a Cas9-mediated transformation. However, double crossover events resulting in a gene deletion rather than disruption were less frequently observed. Therefore, we aimed to enhance the efficiency of gene targeting by deleting the *ku80* (SPBR_02356 and SPSK_07043) gene. The absence of this gene should favour a homologous recombination-mediated repair during the transformation process. To achieve this, we transformed both strains of *S. brasiliensis* with a disruption cassette containing the hygromycin B resistance gene (*hph*), the *PbLuc* luciferase expression construct, and flanking homology arms to the upstream and downstream regions of the *ku80* gene (designed to delete the entire *ku80* coding sequence) (Figure 3A and Supplemental Table 1). The *Paracoccidioides ef1* promoter-*PbLuc* marker was chosen due the high efficiency in generating bioluminescent *Sporothrix* transformants observed during standardization of the protoplast transformation protocol.

**Figure 3:**
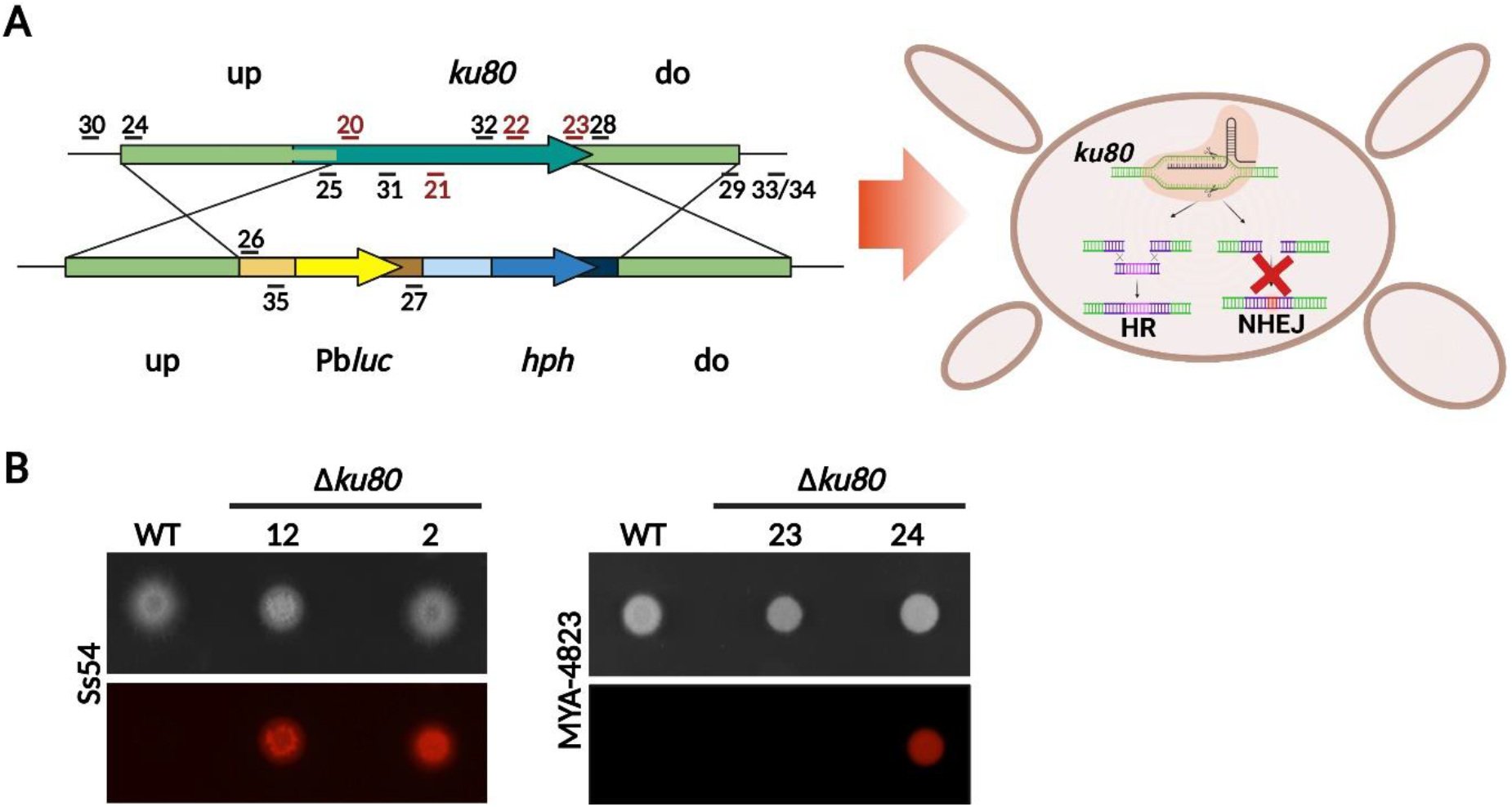
Generation of non-homologous end joining deficient strains of *Sporothrix brasiliensis*. **(A)** Schematic representation of *ku80* deletion cassette containing the *hph* resistance gene, the annealing position of oligonucleotides (black) and gRNAs (red), as well as the schematic representation of the resultant strain defective in NHEJ pathway. **(B)** Bioluminescent imaging of two transformants of Ss54 and MYA-4823 containing in locus integration of the *ku80* deletion cassette inoculated onto GG10_THI_ containing d-luciferin (0.2 mM). Brightfield (upper panels) and luminescence (bottom panels) were recorded after 5 days of incubation at 25 °C.

Putative hygromycin B resistant *ku80^-^* transformants were tested for the presence of the construct by diagnostic PCR. Of the 72 transformants tested for each strain, three Ss54 colonies and two MYA-4823 colonies ultimately showed a single integration of the donor DNA at the expected site when tested by Southern blot analysis (Supplemental Figure 3). Most of these transformants also displayed the expected bioluminescence from luciferase activity (Figure 3B). It is worth noting that, for both strains, the use of a reduced concentration of Cas9 (9.5 pmol) yielded a higher number of transformants compared to the previous *pks1* transformation approaches. Strains with homology-directed repair in either the upstream or downstream regions were also identified in addition to the transformants in which the *ku80* gene had been successfully deleted.

### *ku80* gene deletion enhances targeted DNA integration in *S. brasiliensis*

To test whether the *ku80* deletion enhances targeted integration of the donor DNA during transformation, we repeated the *pks1* deletion using the *S. brasiliensis* strains Ss54_Δ*ku80*_12 and MYA-4823_Δ*ku80*_24. In the presence of Cas9, we obtained 25 transformants from the Ss54_Δ*ku80* transformation, thirteen of which generated white colonies in the first streak on GG10_THI_ supplemented with nourseothricin. The remaining fourteen brown colonies did not survive the subsequent passages for isolation of spores from the transformants. This indicated that, in these fourteen cases, the gene deletion construct had not been stably integrated into the genome. In the MYA-4823_Δ*ku80* transformation, 32 colonies were recovered with 25 forming a white colony and seven showing a brown phenotype. The transformations lacking Cas9 did not generate any white colonies for MYA-4823_Δ*ku80* or Ss54_Δ*ku80*.

Diagnostic PCR analysis was performed on thirteen transformants from each strain to evaluate the effect of *ku80* gene deletion on the accuracy of gene targeting. We observed that accurate targeted deletion of *pks1* had taken place in 100% of the MYA-4823_Δ*ku80* transformants and 85% of the Ss54_Δ*ku80* transformants (Figure 4A). Furthermore, Southern blot analysis of seven MYA-4823_Δ*ku80*_Δ*pks1* transformants and eight Ss54_Δ*ku80*_Δ*pks1* transformants confirmed a single integration of the deletion cassette in all but one of the MYA-4823_Δ*ku80*_Δ*pks1* transformants tested (Figure 4B).

**Figure 4:**
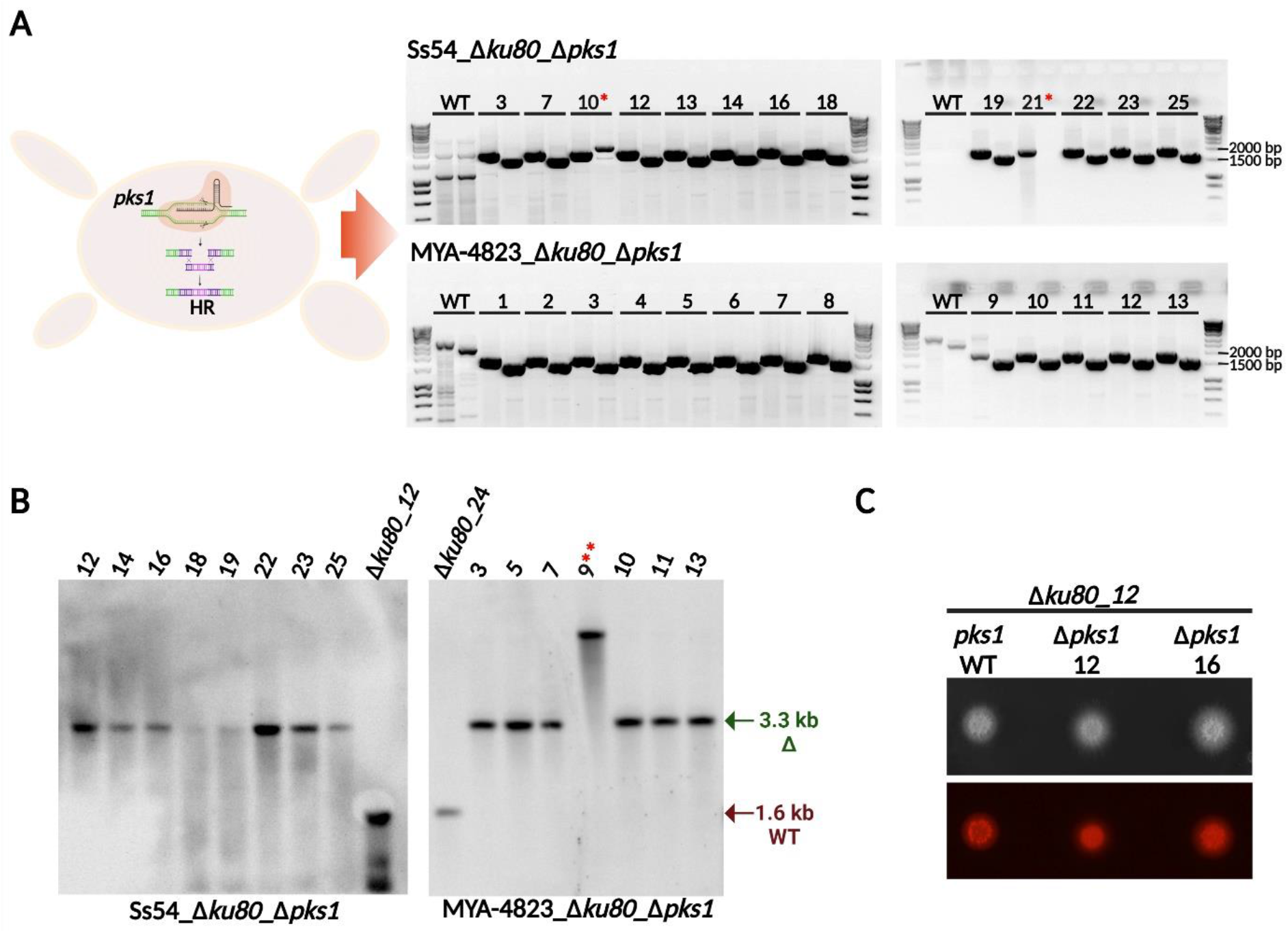
Enhanced homologous recombination by Δ*ku80* strains. **(A)** PCR analysis of putative Δ*ku80*_Δ*pks1* transformants to confirm the disruption of *pks1* gene and in locus integration of the nourseothricin cassette using oligonucleotides for the deletion construct. For each transformant the left-hand lane represents PCR diagnosis of the upstream integration site, and the right-hand lane the diagnosis of the downstream integration site: (*) samples lacking one of the PCR products or containing fragments of unexpected size. **(B)** Southern blot analysis of randomly selected Δ*ku80*_Δ*pks1* transformants from *S. brasiliensis* strains. Diagnostic bands of 1.6 kb for wild type (WT) and 3.3 kb for a deletion mutant (Δ) are highlighted: (**) samples containing a band of unexpected size. **(C)** Bioluminescence imaging of two Ss54_Δ*ku80*_Δ*pks1* transformants and their parental strain Ss54_Δ*ku80*_12; 6×10^5^ spores were spotted onto GG10_THI_ plates containing d-luciferin (0.2 mM). Brightfield (upper panel) and luminescence (bottom panel) were recorded after 5 days of incubation.

To verify the stability of integration of the exogenous DNA across the transformations, spores from two of the Ss54_Δ*ku80*_Δ*pks1* transformants and their parental strains were spotted on GG10_Thi_ supplemented with d-luciferin. Bioluminescence imaging confirmed the stable expression of the luciferase in the Ss54_Δ*ku80*_Δ*pks1* transformants even after this new round of gene deletion (Figure 4C). We concluded that the deletion of *ku80* facilitates the precise integration of the donor DNA during Cas9-mediated transformation, facilitating the screening process for true gene deletion mutants in *S. brasiliensis*.

### *S. brasiliensis* Pks1 mediates oxidative stress protection and resistance during phagocytosis

We tested whether Pks1-dependent melanin production, most likely resulting in DHN-melanin, is responsible for the brown pigmentation of yeasts and mycelia during the long-term cultivation in liquid cultures. Sabouraud and YPD media were inoculated with spores from two independent Ss54_Δ*ku80*_Δ*pks1* mutants and their parental Δ*ku80* and wild-type strains. After 19 days of growth at 25°C, the mycelia from Ss54 WT and Δ*ku80* strains, but not from the Δ*ku80*_Δ*pks1* strains, developed a strong brown pigmentation (Figure 5A). Similarly, Ss54 WT and Δ*ku80* yeasts cultivated in YPD at 37°C for at least seven days showed a darker pigmentation when compared with the Δ*ku80*_Δ*pks1* strains (Figure 5B).

**Figure 5:**
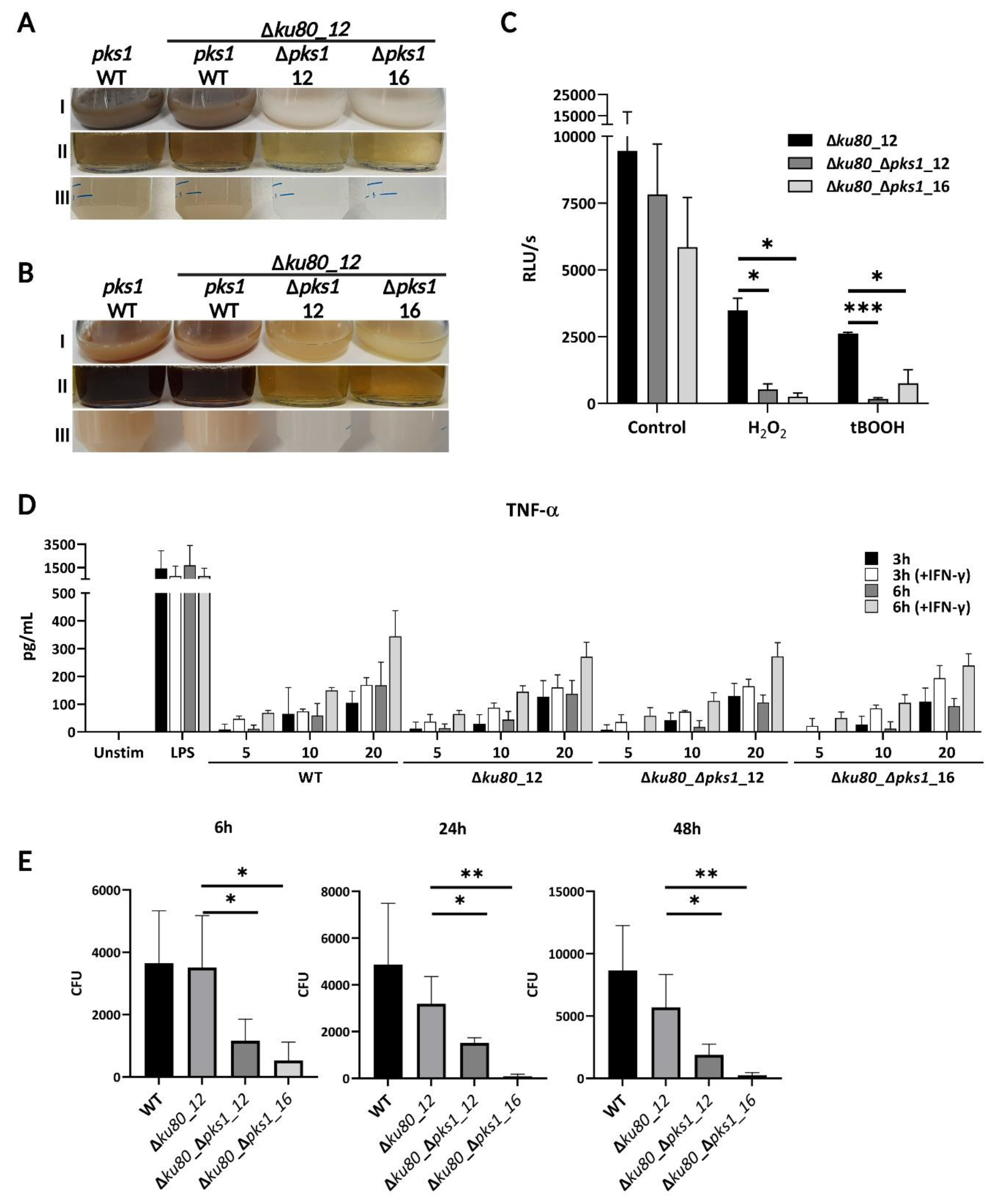
Phenotypic characterization of *Sporothrix* deletion strains. **(A)** The mycelial phase of Ss54 WT, Ss54_Δ*ku80*_12, Ss54_Δ*ku80*_Δ*pks1_12* and Ss54_Δ*ku80*_Δ*pks1*_16 strains was cultivated in Sabouraud dextrose at 25 °C and 150 rpm for 19 days. **(B)** Yeast phase of the same strains cultivated in YPD at 37 °C and 200 rpm for 7 days. For (A) and (B): (I) spores or yeast cultures; (II) culture supernatants; and (III) suspensions of washed spores or yeast cells. **(C)** Bioluminescence assay of yeast cells cultured in RPMI containing 0.4 mM d-luciferin and exposed to 20 mM hydrogen peroxide (H_2_O_2_) or 8 mM tert-butylhydroperoxide (tBOOH) for 8h. Each assay was performed in technical triplicate and repeated twice, the statistical differences were calculated by mean values using student’s t-test: *, p< 0.05; ***, p< 0.0005. **(D)** TNF-α levels after 3 or 6 h of co-incubation of BMDM with opsonised yeast cells from Ss54 WT and *pks1* deletion strains. BMDMs were either primed or not primed with INF-γ and were co-incubated with yeast cells at an MOI of 5, 10 or 20. **(E)** Phagocytosis of Ss54 WT and *pks1* deletion strains by BMDM. Opsonised yeast cells were co-incubated with BMDM that had been primed with INF-γ for 2 h. Non-phagocyted *Sporothrix* cells were washed off after 3 h of co-incubation and CFUs were evaluated after additional 3 (6 h), 21 (24 h) and 45 (48h) hours of incubation with the primed BMDM. Statistical differences were calculated using student’s t-test: *, p < 0.05; **, p < 0.01.

The production of melanin is a virulence determinant due to the ability of melanin to scavenge free radicals. This confers the resistance of fungal pathogens to host-pathogen interactions, as well as to environmental stresses (Eisenman and Casadevall, 2012). A previous study had suggested that inhibition of the DHN-melanin pathway results in impaired survival of *S. schenckii* to oxidative stress induced by hydrogen peroxide (H_2_O_2_) and phagocytosis (Romero-Martinez *et al*., 2000). Therefore, to test whether *pks1* deficient strains are more sensitive to oxidative stress, we measured the light emission from the luciferase activity of live cells after exposure to H_2_O_2_ or tert-butylhydroperoxide (tBOOH). In comparison to the parental strain, exposure to 20 mM H_2_O_2_ significantly decreased the intensity of the bioluminescence generated by both of the Ss54_Δ*ku80*_Δ*pks1* mutants (Figure 5C). Similarly, the bioluminescence signal intensity was significantly higher from the parental strain Ss54_Δ*ku80* than from the Δ*pks1* mutants after exposure to 8 mM tBOOH. This suggests that the presence of a functional *pks1* gene confers protection against the oxidative stresses imposed by tBOOH and H_2_O_2_.

To test whether Pks1 influences fungus-host interactions, bone marrow-derived macrophages (BMDM) were primed or not with interferon-γ (INF-γ) and co-incubated with opsonised yeasts of Ss54 WT, Ss54_Δ*ku80*_12 and two Ss54_Δ*ku80*_Δ*pks1* mutants for 3 and 6 h at a multiplicity of infection (MOI) of 5, 10 and 20. Tumour necrosis factor alpha (TNF-α) was measured from the supernatants. TNF-α secretion by BMDMs was proportionate to the MOI and priming of the cells with IFN-γ resulted in a slight but non-significant increase of TNF-α secretion. Neither deletion of *ku80* nor *ku80* plus *pks1* resulted in a modified secretion of TNF-α by BMDMs compared to the response induced by Ss54 WT yeasts (Figure 5D).

To compare the resistance of Ss54 WT, Ss54_Δ*ku80* and Ss54_Δ*ku80*_Δ*pks1* cells to phagocytic killing, the fungal strains were opsonised with mouse serum and co-incubated with INF-γ-primed BMDMs. After 3 h of co-incubation, the extracellular cells of *Sporothrix* were removed by washing and the colony-forming units (CFU) were determined after additional 3 (total of 6 h), 21 (total of 24 h) and 45 (total of 48 h) hours of incubation (Figure 6C). The two independent Ss54_Δ*ku80*_Δ*pks1* transformants revealed a statistically significant reduction in CFU counts compared to their parental strain. Therefore, we conclude that inactivation of *pks1* significantly affects survival of *Sporothrix* cells after phagocytosis and plays an important role in host-pathogen interactions.

## DISCUSSION

Genetic engineering is a key tool for dissecting the molecular mechanisms that underlie virulence and for facilitating the search for more efficient antimicrobial therapies. *Sporothrix* species have been considered challenging for genetic manipulation (Mora-Montes *et al*., 2015) and only limited tools have been available for applying reverse genetics to resolve molecular mechanisms critical for sporotrichosis development (Lozoya-Perez et al., 2019; Lozoya-Perez *et al*., 2018; Romero-Martinez *et al*., 2000; Song *et al*., 2021; Tamez-Castrellon *et al*., 2018; Tamez-Castrellon *et al*., 2021). RNA interference-mediated gene silencing using *Agrobacterium*-mediated transformation has been used for studying the Golgi α-1,6-mannosyltransferase (*Och1*) and protein rhamnosylation (*rmlD*) genes in *S. schenckii* (Lozoya-Perez *et al*., 2019; Tamez-Castrellon *et al*., 2021). However, so far, attempts to generate platform strains to facilitate the screening of transformants and to monitor the effects of gene deletion or overexpression using *Sporothrix* as model organism have not been reported.

In this study, we have successfully established an efficient protocol for the accurate integration of exogenous DNA into the genomes of *S. brasiliensis*, *S. chilensis* and *S. schenckii*. We applied PEG-mediated fusion of protoplasts that were generated from the mycelial phase to stably incorporate a plasmid containing *hph* and *PbLuc* genes into the *Sporothrix* genome. Light emission from the luciferase reporter was still detectable after a vegetative sub-cultivation period of three years, which is consistent with the stable integration of genes into the genomes of *Sporothrix* species. In addition, the functional expression of a red-shifted luciferase enables real-time monitoring of *in vivo* of disease progression as shown for other fungal species such as *Aspergillus fumigatus*, *Aspergillus terreus*, *Candida albicans* and *Cryptococcus neoformans* (Brock et al., 2008; Galiger et al., 2013; Jacobsen et al., 2014; Poelmans et al., 2018; Resendiz-Sharpe et al., 2022; Seldeslachts et al., 2021; Slesiona et al., 2012; Vande Velde et al., 2018; Vanherp et al., 2019).

For targeted gene disruption and deletion, previous studies have reported that flanking homologies ranging in length from 30 to 2000 bp can yield efficient homology-directed repair after DNA double-strand breaks induced by Cas9 in aspergilli, *Rhizopus microspores* and *C. neoformans* (Huang et al., 2022; Lax et al., 2021; Nodvig et al., 2018). Therefore, based on the protocols used for gene deletion in *Magnaporthe oryzae* and *Aspergillus* species (Foster et al., 2018; Peres da Silva and Brock, 2022), we developed a CRISPR/Cas9-based protocol for gene disruption in *S. brasiliensis* and *S. schenckii*. By exploiting the putative 1,3,6,8-tetrahydroxynaphtalene synthase gene *pks1* as a visual marker for successful gene disruption/deletion and resistance to nourseothricin as transformation marker, we confirmed that DNA double-strand breaks were successfully induced by Cas9 to generate mutants deficient for DHN-melanin like pigments. However, even though Cas9 precisely cut at the expected *pks1* Cas9 sites, homologous recombination at both flanks was rarely observed in wild-type *Sporothrix* backgrounds. This suggested a low rate of homology-directed DNA repair in *S. brasiliensis* and *S. schenckii* during the transformation process, as described for other fungi (Choquer et al., 2008; da Silva Ferreira et al., 2006; Goins et al., 2006).

The repair of DNA double-strand breaks is assumed to occur through two main pathways: homologous recombination and non-homologous end joining repair (NHEJ). The NHEJ is the most active pathway for DNA repair in filamentous fungi, which often reduces the efficacy of in locus DNA integration during the transformation process (Krappmann, 2007). The Ku70/Ku80 heterodimer participates in the initial step of DNA double-strand repair by the NHEJ pathway. For this reason, the deletion of either *ku70* or *ku80* encoding genes has been used successfully as a strategy to improve homologous recombination in a diverse set of fungal species (Choquer *et al*., 2008; Goins *et al*., 2006; Peres da Silva and Brock, 2022; Resendiz-Sharpe *et al*., 2022). The deletion of the *ku80* gene in the Ss54 and MYA-4823 strains of *S. brasiliensis* improved the efficiency of homologous recombination during the transformation to an index greater than 80%. The Δ*ku80* transformants preserved the expected wild-type morphology, as well as their thermo-dimorphism.

In terms of the successful deletion of the *pks1* gene, we observed a change in colony and medium colouration for *pks1* mutants compared to the wild-type parental strain, which is compatible with the lack of production of pigments like DHN-melanin. Increased melanin production is observed during responses to cellular stresses such as starvation, oxidative and nitrosative stresses (Lee *et al*., 2019; Missall *et al*., 2005; Poyntner *et al*., 2018). Accordingly, deletion of enzymes related to melanin biosynthesis results in attenuated fungal virulence (Akoumianaki *et al*., 2016; Lee *et al*., 2019; Li et al., 2022; Missall *et al*., 2005; Poyntner *et al*., 2018; Xiao et al., 2021). In this respect, it has been shown that DHN-melanin from *Aspergillus fumigatus* disrupts the phagosomal recruitment of Ca^2+^/calmodulin-dependent protein kinase and the autophagy pathway LC3-associated phagocytosis preventing the elimination of conidia (Akoumianaki *et al*., 2016; Goncalves et al., 2020). Previous studies have shown that *S. schenckii* and *S. globosa* UV-induced mutants deficient for DHN-melanin production are more susceptible to oxidative/nitrosative stresses and induce higher levels of reactive oxygen species by human monocytes and murine macrophages (Guan *et al*., 2021; Romero-Martinez *et al*., 2000). Furthermore, a *S. globosa* DHN-melanin deficient strain is less resistant to phagocytosis (Guan *et al*., 2021; Masternak *et al*., 2000). Therefore, the sensitivity of *pks1* mutants to H_2_O_2_ and tBOOH observed in this study was consistent with the view that pigments like DHN-melanin protect *Sporothrix* against oxidative stress. The absence of pigments like DHN-melanin probably also accounts for the elevated sensitivity to phagocytic killing by BMDM. Most importantly, our targeted deletion of *pks1* provides a proof-of-concept for future investigations of selected genes and pathways in *Sporothrix*.

In summary, our results show that PEG- and CRISPR/Cas9-mediated transformation of protoplasts provides a precise and efficient tool for the genetic manipulation of pathogenic *Sporothrix* species. Furthermore, efficient gene targeting by homologous recombination is significantly enhanced in strains deficient in NHEJ repair. Our developments on genetic tools to study *Sporothrix* species complex been reinforced by a parallel study that also describes approaches for heterologous gene expression in *S. brasiliensis* (see adjoining paper by Tavares and coworkers 2022). The development of reliable and reproducible tools for genetic engineering of *Sporothrix* opens up powerful experimental avenues to improve the understanding of mechanisms underlying the development of sporotrichosis. This is particularly important given the rapid geographic expansion and upsurge of atypical and more severe forms of this debilitating fungal disease.

## ACKNOWLEDGMENTS

We are grateful to Professor Anderson Messias Rodrigues (Federal University of São Paulo-Brazil) for providing strains of *S. brasiliensis* and *S. schenckii*.

## FUNDING

RPS was financially supported by a fellowship from the Medical Research Council Centre for Medical Mycology at University of Exeter (MR/N006364/2). MB was supported by the Medical Research Council (MR/N017528/1). AJPB was supported by a programme grant from the UK Medical Research Council (www.mrc.ac.uk: MR/M026663/1, MR/M026663/2). GDB was supported by the Wellcome Trust (102705, 217163).

## AUTHORS CONTRIBUTION

RPS conceived the study, performed experiments, analysed the data and wrote the first draft of the manuscript. RH performed BMDM stimulation assay and cytokine measurements. IL performed oxidative stress assays. MB, AJPB and GDB contributed to the design of the study, interpretation of data and the writing and critical reviewing of the manuscript. All authors reviewed and approved the final version of the manuscript.

## DECLARATION OF INTERESTS

The authors declare no conflict of interest.

## METHODS

### Strains and growth conditions

*Sporothrix brasiliensis* Ss54, *S. schenckii* Ss126 and *S. chilensis* Ss469 strains were kindly provided by Professor Anderson Messias Rodrigues (Federal University of São Paulo) and the *S. brasiliensis* MYA-4823 strain was acquired from ATCC.

For harvesting spores, strains were cultivated for 7 days in Sabouraud liquid medium (Sigma) pH 4.5 at 25 °C and 150 rpm. Cultures were filtered through Miracloth (Merck) filter gauze, supernatants centrifuged at 3220 x*g* for 10 min and spores washed twice in phosphate-buffered saline (PBS; Gibco). Fungal transformants were plated on GG10 medium (Geib and Brock, 2017; Geib et al., 2016) containing 3 μg/ml thiamine (Sigma) (GG10_THI_) supplemented with 1.2 M sorbitol (Sigma), 80 μg/ml of nourseothricin (nat, Jena Biosciences) or 180 μg/mL of hygromycin B (Thermo) when necessary. Yeast cells were grown at 37 °C in Yeast Peptone Dextrose (YPD; 1% yeast extract-Difco, 2% bacteriological peptone-Oxid, 2% glucose-Fisher) pH 7.8, liquid cultures were kept at 200 rpm and solid YPD was supplemented with 2% (w/v) of agar. For cytokine stimulation, phagocytic killing and oxidative stress assays, spore suspensions were inoculated in RPMI medium (Sigma) supplemented with 2% (w/v) glucose and non-essential amino acids (Gibco), pH 7.2 (abbreviated as RPMI 2% glucose hereafter) and incubated at 37 °C and 200 rpm for 6 and 7 days, respectively.

### Kits and reagents

DNA amplifications for cloning and transformant screening were performed with Phusion Green High-Fidelity DNA Polymerase (Thermo) or Phire Green Hot Start II DNA Polymerase (Thermo), respectively. For cloning, the PCR products and digested plasmids were run on 1 or 0.8% agarose gels, and gel elution of DNA fragments was performed using Zymoclean Gel DNA Recovery Kit (Zymo). The *in vitro* assembly of PCR products was performed by using In-Fusion HD cloning kit (Takara). The assembled plasmids were amplified by transformation of Stellar Competent Cells (Takara) and purified from overnight cultures in LB medium containing 100 μg/ml Ampicillin grown at 37°C by using the NucleoSpin Plasmid isolation kit (Macherey-Nagel). The selection markers for fungal transformation were cloned into digested plasmids by using the Rapid DNA Ligation Kit (Thermo). The guide RNAs (gRNAs) were generated by EnGen sgRNA Synthesis Kit (NEB) according to the manufacturer’s protocol. The synthesized gRNA was purified by RNA Clean & Concentrator (Zymo) and assembled to the EnGen Spy Cas9 NLS (NEB) directly before the fungal transformation. Oligonucleotide sequences utilized in this study are described in the Supplementary Table 1.

### Transformation constructs

To establish the protocol of transformation for *Sporothrix*, we first tested plasmids that were previously used in *Aspergillus niger* and/ or *Paracoccidoides* transformations (Milhomem Cruz-Leite *et al*., 2022). We also generated the P*ef1:Pbluc_OPT_red_*:T*eno1*_URABlaster_pUC19 by amplification and assembly of the synthetic gene *Pbluc_OPT_red_*(GenBank: MT978127), as well as the *elongation factor 1-gamma* (*ef1*, PAAG_03556) promoter and *enolase 1* (*eno1*, PAAG_11169) terminator from Pb01 genomic DNA (gDNA), using the oligonucleotides from 1 to 6. To replace the URABlaster selection marker, P*act:Pbluc_OPT_red_*:T*eno1*_URABlaster_pUC19, P*eno1:Pbluc*:T*eno1*_URABlaster_pUC19 (Milhomem Cruz-Leite *et al*., 2022) and P*ef1:Pbluc_OPT_red_*:T*eno1*_URABlaster_pUC19 (this study) were digested with *Not*I (Thermo) and Alkaline Phosphatase (AP; Thermo), gel purified and assembled with a *Not*I-restricted hygromycin B (*hph*) resistance cassette.

The constructs for the *pks1* (SPSK_00653; SPBR_06313) deletion cassettes were based on the sequence identified in *C. lagenarium pks1* gene (Fujii *et al*., 1999) and the *A. fumigatus* naphthopyrone synthase *pksP* gene (Langfelder *et al*., 1998; Watanabe *et al*., 2000). A sequence of 892 bp upstream of *pks1* gene, including the first 17 bp of the coding region, was amplified by PCR using the oligonucleotides 9 and 10 and MYA-4823 gDNA as template. The 862 bp downstream region of the *pks1* deletion construct was designed starting from the 2233 bp (*S. brasiliensis*) or 2229 bp (*S. schenckii*) of the *pks1* using the oligonucleotides 11 and 12. In the reverse oligonucleotide for the upstream flanking region, a stop codon sequence was included as well as a *Not*I restriction site for cloning of the selection marker. The upstream and downstream flanking fragments of the *pks1* deletion construct were assembled by *in vitro* recombination into a *Sma*I-restricted pUC19 plasmid (Thermo)*. E. coli* transformants containing the correct plasmid were selected by colony PCR using oligonucleotides 12 and 13. The isolated Δ*pks1*_pUC19 plasmids were digested with *Not*I plus AP and assembled with the *nat1* resistance cassette *Not*I-digested. The correct assembly was confirmed by colony PCR using the oligonucleotides 12 and 19. For *Sporothrix* transformation, the deletion cassette Δ*pks1::nat1* was excised from pUC19 by digestion with *Sma*I. The two gRNAs for *pks1* deletion were synthetised using the oligonucleotides 7 and 8.

The ATP-dependent DNA helicase 2 subunit 2 (*ku80*) from *Sporothrix* species (SPBR_02356 and SPSK_07043) were identified by Blast search using the *Aspergillus fumigatus akuB* sequence (AFUA_2G02620) as template at NCBI or EnsemblFungi websites (https://www.ncbi.nlm.nih.gov/, https://fungi.ensembl.org/index.html). The upstream region of the construct (774 bp) including 411 bp of the coding region as well as the downstream region (792 bp) were amplified from Ss126 gDNA using the oligonucleotides 24 and 25 or 28 and 29, respectively. The *Pef1:Pbluc_OPT_red_:Teno1* fragment was amplified from the plasmid described above using oligonucleotides 26 and 27. The three PCR products were *in vitro* assembled in a *Sma*I digested pUC19 plasmid. The Δ*ku80::Pbluc*_*OPT_red*__pUC19 plasmid was digested by *Not*I plus AP and assembled with the *Not*I-restricted *hph* selection marker. For *Sporothrix* transformation, the deletion cassette Δ*ku80::Pbluc*_*OPT_red*__*hph* was released from the pUC19 backbone by digestion with *Sma*I. The four gRNAs utilized for the *ku80* deletion were synthetised using the oligonucleotides 20, 21, 22 and 23.

### Transformation

*Sporothrix* spore suspensions were inoculated in 50 ml of Sabouraud pH 4.5 and cultivated for 48 to 72 h at 25 °C and 150 rpm. Protoplasts were prepared as described previously for *Aspergillus* species (Peres da Silva and Brock, 2022) with some modifications. The mycelium was collected and washed with sterile tap water over sterile Miracloth filter gauze and transferred into 30 ml of 90 mM citrate/phosphate buffer pH 7.3 supplemented with 10 mM 1,4-Dithiothreitol (DTT, Thermo) and incubated for 1 h at 25°C. The mycelium was collected and washed with citrate/phosphate buffer over sterile Miracloth and transferred to 20 ml of protoplasting solution (0.6M (NH_4_)_2_SO_4_, 50 mM maleic acid buffer pH 5.5) containing 100 mg Yatalase (Takara) and 100 mg Lysing Enzymes from *Trichoderma harzianum* (Sigma) and was incubated at 25°C and 70 rpm whereby the release of protoplasts was monitored under the microscope for up to 90 min. The protoplasts were separated from the mycelium by filtration over sterile Miracloth, pelleted at 3220 x*g* for 8 min at 4 °C, washed once in 20 ml of washing solution (0.6 M KCl, 0.1 M Tris/HCl, pH 7.0), followed by resuspension in 10 ml of solution A (50 mM CaCl_2_, 0.6 M KCl, 0.1 M Tris/HCl pH 7.5). The number of protoplasts was evaluated using a Neubauer counting chamber and the protoplasts were finally resuspend in solution A to give a concentration of about 5 ×10^6^ to 2 × 10^7^ protoplast/ml and 100 μl of this suspension was used in the transformation procedure.

The uptake of donor DNA and Cas9 by protoplasts was adapted from protocols previously described for *Magnaporthe oryzae* and *Aspergillus species* (Brock *et al*., 2008; Foster *et al*., 2018; Gressler et al., 2011; Peres da Silva and Brock, 2022). The Cas9/gRNA complex was generated as described by the manufacturer’s protocol (NEB). For *pks1* deletion, a total of 80 pmol of each synthetised gRNA was mixed with 40 pmol of Cas9, in two different microtubes, and incubated for 10 min at 25 °C before adding the complex to the protoplasts. 1 μg of deletion cassette was added to a 2 ml microtube together with the two Cas9/gRNA complex. In transformations performed only in the presence of the donor DNA, 1.5 μg of the deletion cassette was added to each microtube. For targeting the *ku80* gene, the gene deletion was performed with 9.5 pmol Cas9 per 19 pmol gRNA for each of the gRNAs utilized. 1 μg or 1.5 μg of deletion cassette were added to 2 ml microtubes with or without Cas9/gRNA complexes, respectively. The 100 μl of protoplast suspension were mixed with 25 μl of PEG solution (25% PEG 8000, 50 mM CaCl_2_, 10 mM Tris/HCl, pH 7.5) in a 2 ml microtube containing Cas9/gRNA complexes and the donor DNA or only the later. The microtube was incubated for 25 min at room temperature and another 500 μl of PEG solution was added. After 5 min, 1 ml of solution A was added and the protoplast suspension transferred to 5 mL of Sabouraud supplemented with 1.2 M sorbitol, pH 4.5. For regeneration, the protoplasts were incubated overnight at 25°C at 100 rpm, pelleted at 3220 x*g* for 8 min at room temperature, washed twice in 5 ml of solution A and resuspended in 1.6 ml of solution A. 400, 500 and 700 μl of the resuspended regenerated protoplast were plated in three different agar plates of GG10_THI_ supplemented with 1,2 M Sorbitol and 80 μg/mL of nourseothricin or 180 μg/mL of hygromycin B.

### Molecular analysis of transformants

After three passages in GG10_THI_ supplemented with nourseothricin or hygromycin B, spores from selected transformants were transferred to a 48-well plate containing 500 μl of YPD and incubated for 5 days at 37 °C and 180 rpm. The cells were harvested in a screw-cap tubes containing Lysing Matrix Y (MP Biomedicals) and pelleted at 10000 x*g* for 10 min at 4 °C. The supernatant was discarded, 500 μl of extraction buffer (100 mM Tris/HCl, 50 mM EDTA, 500 mM NaCl, 10 mM β-mercaptoethanol, 1% SDS, 20 μg RNAse A) was added and the cells were disrupted in a FastPrep machine (MP Biomedicals) in three rounds of 30 s at 6000 rpm. For Southern blot analysis, protoplasts were prepared as described in the transformation procedure, but 1.3 g of Vino Taste Pro (Novozyme) in osmotic solution (0.6 M KCl, 10 mM phosphate buffer pH 5.7) were utilized instead of the yatalase enzyme. After overnight incubation at 25 and 70 rpm, the protoplasts were separated from the mycelium by filtration over Miracloth, pelleted at 3220 x*g* for 8 min at 4 °C, resuspended in 500 μl of extraction buffer and transferred to 2 ml microtubes. The gDNA from disrupted yeast cells and from protoplast was extracted as described by Dellaporta *et al*. (1983) (Dellaporta et al., 1983). The gDNA from protoplasts preparations was purified from 0.8 % agarose gel using Large Fragment DNA Recovery Kit (Zymo).

The screening of the *pks1* deletion mutants was performed by PCR to verify the absence of the respective coding region (oligonucleotides 14 and 15 or 16 and 17) and the in locus insertion of the deletion constructs (oligonucleotides 14 and 18 or 17 and 19). The *ku80* deletion strains were similarly screened for the excision of the whole coding region (oligonucleotides 30 and 31 or 32 and 33/34) and the in locus insertion of *luciferase* and *hph* genes (30 and 35, 26 and 33/34 or 30 and 33/34). In addition, selected Δ*ku80* and Δ*ku80*_Δ*pks1* strains were plated on Sabouraud and YPD agar paltes supplemented with 0.4 mM of d-luciferin potassium salt (Promega) to confirm the expression of a functional luciferase by bioluminescence imaging using ChemiDoc MP (Bio-Rad).

### Southern blot analysis

To confirm the insertion of the deletion construct into the genome by homologous recombination, gel-purified gDNA from potential Δ*pks1* and Δ*ku80*_Δ*pks1* transformants and their parental strains were digested overnight by *Pst*I (Thermo). gDNA of Δ*ku80* strains was digested overnight with *Aat*II (Thermo). Digested gDNA was loaded on a 0.8% agarose gel, separated by electrophoresis and transferred to a nylon membrane Hybond N+ (GE Healthcare). The probes for the upstream regions of the *pks1* (oligonucleotides 9 and 10) and *ku80* (oligonucleotides 24 and 25) deletion constructs were synthetised using PCR DIG DNA Labeling Mix (Roche) and Taq DNA Polymerase (NEB). Probes were hybridized overnight in DIG Easy Hyb (Roche) at 42 °C and detection was performed by chemiluminescence using anti-Digoxigenin-AP Fab fragments (Roche) and CDP-Star chemiluminescent substrate (Roche) in a ChemiDoc MP (Bio-Rad).

### Oxidative stress assay

Yeast cells of selected strains were cultivated in RPMI 2% glucose, pelleted at 3220 x*g* for 10 min at 4 °C, washed twice in PBS and resuspended at a concentration of 1.25 × 10^7^/ml in fresh RPMI 1640 GlutaMAX-I 25 mM HEPES (Gibco) containing 0.4 mM d-luciferin without addition of an oxidative stressor (control) or supplemented with 20 mM hydrogen peroxide (H_2_O_2_; Sigma) or 8 mM tert-butylhydroperoxide (tBOOH; Sigma). The yeasts were incubated at 37 °C and 5% CO_2_. The relative light units per second (RLU/s) were measured after 8 h using a TECAN Spark plate reader set at 37 °C and 5% CO_2_ in the air space. The assays were performed at least 2 times in triplicates. The mean of luminescence and standard deviations were plotted. Statistically significant differences were calculated using student’s t-test in GraphPad Prism version 9 for Windows.

### Isolation of BMDMs

For isolation of murine bone-marrow from femurs and tibias, we have used four 6-8 weeks old male and female C57BL/6 mice bred in-house at the University of Exeter and housed in stock cages under pathogen-free conditions.

Bone-marrow-derived macrophages were obtained as described by others (Davies and Gordon, 2005). Briefly, bone-marrow was extracted from femurs and tibias of C57B6 mice and cultured in RPMI 1640 GlutaMAX-I 25 mM HEPES, 100 μg/mL penicillin/streptomycin (abbreviated as RPMI hereafter), supplemented with 10% (v/v) Fetal Bovine Serum (FBS) and 20% (v/v) L929-conditioned supernatant on day 1. Medium was replaced on day 4 and cells were used for subsequent experiments on day 7.

### BMDM stimulation and quantification of cytokine production

BMDMs were brought into suspension with PBS containing 8 mg/mL lidocaine and 5 mM EDTA, counted and seeded at 2 × 10^5^ cells per well in 24-well plates in RPMI medium supplemented with 10% FBS. After overnight incubation at 37 °C and 5% CO_2_, the medium was replaced with RPMI or RPMI containing 50 ng/mL IFN-γ (Abcam). After 1 h at 37 °C and 5% CO_2_, the supernatant was discarded and replaced with fresh RPMI without FBS. Yeasts cultivated in RPMI 2% glucose were centrifuged at 3220 × g for 5 min, washed three times with PBS (Gibco) and counted in haemocytometer chamber. 2 × 10^7^ yeasts were opsonised in 1 mL RPMI 10% (v/v) mouse serum (Sigma) at 37 °C for 1 h, then centrifuged, washed once in PBS and resuspended in serum-free RPMI. For the cytokine assays, the BMDM with and without pre-stimulation with INF-γ were co-incubated with opsonised yeasts at the indicated multiplicity of infection (MOI) or with LPS (10 ng/mL) for 3 or 6 h. Supernatant was collected, centrifuged at 4,000 × *g* for 5 min to remove remaining yeast cells and the yeast-free supernatants were kept frozen until cytokine ELISA was performed. The concentration of TNF-α was determined using the DuoSet ELISA kit (R&D systems) according to the manufacturer’s protocol. The statistical analyses were performed using student’s t-test and GraphPad Prism version 9 for Windows.

### *Sporothrix* survival assay

BMDMs were seeded at 2 × 10^5^ cells per well in 48-well plates in RPMI medium containing 10% FBS. After overnight incubation, the cells were primed with 40ng/mL in RPMI medium containing 10% FBS for 2 h at 37 °C and 5% CO_2_. Yeasts cultivated for 7 days in RPMI 2% glucose were pelleted and opsonised as described above. Opsonised yeasts at MOI 5 were co-incubated for 3 h with BMDM IFN-γ activated at 37 °C and 5% CO_2_. Three washes with PBS were performed by pipetting up and down to remove extracellular yeasts and the medium was replaced by fresh RPMI 10% FBS. BMDMs with firmly attached or internalized yeasts were incubated for additional 3 h (total = 6h), 21 h (total = 24 h) or 45 h (total = 48 h) at 37 °C and 5% CO_2_. After three washes with PBS, the cells were lysed with 0.01% Triton x-100 in water and 50 μl of undiluted BMDM lysate, as well as 1:10 and 1:100 dilutions were streaked on YPD agar plates. The CFUs were evaluated after 10 to 14 days of incubation at 37 °C. The survival assay was performed in technical duplicate and repeated at least twice for Ss54 WT, Δ*ku80*_12, Δ*ku80*_Δ*pks1*_12 and Δ*ku80*_Δ*pks1*_16. The statistical analyses were performed using student’s t-test and GraphPad Prism version 9 for Windows.

### Figures

All figures were created using BioRender.com.

## Notes

### Competing Interest Statement

The authors have declared no competing interest.

