## Supplementary Material for "CRISPR-based tools for genetic manipulation in pathogenic *Sporothrix* species"

### **Supplementary Table 1:** Oligonucleotides

| Number | Sequence | Target | Protocol |
|---|---|---|---|
| **1** | CTAGAGTCGACCTGCAGCTCGTGCATGTGAAGTTTTCCG | *P. brasiliensis* | Cloning of EF1 promoter, Luc Pb and tENO1 in URA Blaster pUC19 |
| **2** | TCTTGGCATCCTCCATGTTGAAGAACACAGAACGAATAG | *P. brasiliensis* | Cloning of EF1 promoter, Luc Pb and tENO1 in URA Blaster pUC19 |
| **3** | ATGGAGGATGCCAAGAACATCAAG | *PbLuc* | Cloning of Pb promoter, Luc Pb and tENO1 in URA Blaster pUC19 |
| **4** | TCCACGCCCCTATACGGCGATCTTGCCGC | *PbLuc* | Cloning of Pb promoter, Luc Pb and tENO1 in URA Blaster pUC19 |
| **5** | GCCGTATAGGGGCGTGGAGATGAGATGAG | *P. brasiliensis* | Cloning of Pb promoter, Luc Pb and tENO1 in URA Blaster pUC19 |
| **6** | GCTTGCATGCCTGCAGCCACTGATGTTGGAGGTGACT | *P. brasiliensis* | Cloning of Pb promoter, Luc Pb and tENO1 in URA Blaster pUC19 |
| **7** | TTCTAATACGACTCACTATAGCGAGGGACTGGTCTCCAAAGGTTTTAGAGCTAGA | *Sporothrix* | gRNA synthesis for *pks1* deletion |
| **8** | TTCTAATACGACTCACTATAGCTACGTCGAGATGCACGGCAGTTTTAGAGCTAGA | *Sporothrix* | gRNA synthesis for *pks1* deletion |
| **9** | TGAATTCGAGCTCGGTACCCGGGATGTGTGATGTCTGTCTG | *S. brasiliensis* | Deletion contruct *pks1* |
| **10** | GCGGCCGCTTATTCTACTTAGCCAGCCTGTCAGCCATG | *S. brasiliensis* | Deletion construct *pks1* |
| **11** | TAAGTAGAATAAGCGGCCGCGACGGTACCGAGATGCTGTC | *S. brasiliensis* | Deletion construct *pks1* |
| **12** | GTCGACTCTAGAGGATCCCCGGGCCATTTCGAAGCACGC | *S. brasiliensis* | Deletion construct *pks1* |
| **13** | CGTCTTCACTCCAGTCTGTC | *S. brasiliensis* | Control for deletion construct *pks1* |
| **14** | GCCTGTCTCCATTGCTTCG | *Sporothrix* | Deletion control of *pks1* |
| **15** | GAT GCC GTT GGC ACT GGA TG | *Sporothrix* | cloning putative DHN-melanin PKS |
| **16** | GCG CCA AGA CAG AGC TCA C | *Sporothrix* | Deletion control of *pks1* |
| **17** | CCA GCG TCT CAA AGT CGT C | *Sporothrix* | Deletion control of *pks1* |
| **18** | CGG TAT CGG TAG GCG GTG | *nat1* | Deletion control of *pks1* |
| **19** | GGGTTTCACCCTCTGTGGTC | *nat1* | Deletion control of *pks1* |
| **20** | TTCTAATACGACTCACTATAGGCGTCGCATTGCGGGCAAGAGTTTTAGAGCTAGA | *Sporothrix* | gRNA synthesis for *ku80* deletion |
| **21** | TTCTAATACGACTCACTATAGTCTCGAGCTGCGTAATGTTGGTTTTAGAGCTAGA | *Sporothrix* | gRNA synthesis for *ku80* deletion |
| **22** | TTCTAATACGACTCACTATAGACAGATGGGCCAGATTGTGGGTTTTAGAGCTAGA | *Sporothrix* | gRNA synthesis for *ku80* deletion |
| **23** | TTCTAATACGACTCACTATAGTCGGCGCCTAGCACCAAAGAGTTTTAGAGCTAGA | *Sporothrix* | gRNA synthesis for *ku80* deletion |
| **24** | GAATTCGAGCTCGGTACCCGGGTTCATTGCTCAGTATGGTACAG | *S. schenckii* | Deletion construct of *Sporothrix ku80* |
| **25** | TTCACATGCACGAGAGGCCTCGTCCGTCACAAGCACAATC | *S. schenckii* | Deletion construct of *Sporothrix ku80* |
| **26** | GCTTGTGACGGACGAGGCCTCTCGTGCATGTGAAGTTTTCCG | *P. brasiliensis* | Deletion construct of *Sporothrix ku80* |
| **27** | CGGGTCGCGGCCGCAGGCCTCCACTGATGTTGGAGGTGACT | *P. brasiliensis* | Deletion construct of *Sporothrix ku80* |
| **28** | CAGTGGAGGCCTGCGGCCGCGACCCGAGTCAGACATGCG | *S. schenckii* | Deletion construct of *Sporothrix ku80* |
| **29** | GTCGACTCTAGAGGATCCCCGGGCCATGAGCAACCCG | *S. schenckii* | Deletion construct of *Sporothrix ku80* |
| **30** | GATGTCCAATTGCTCACTCTAC | *Sporothrix* | Deletion control of *Sporothrix ku80* |
| **31** | CAT CGA AGT CGA CAC CGC TG | *Sporothrix* | Deletion control of *Sporothrix ku80* |
| **32** | CGGTCCTACCATCAAGAAGG | *Sporothrix* | Deletion control of *Sporothrix ku80* |
| **33** | CAG ATA ACA ACT TGC TGT CCA G | *S. brasiliensis* | Deletion control of *Sporothrix ku80* |
| **34** | GCT AGC AGA TAT TGA CTT GTT GTC | *S. schenckii* | Deletion control of *Sporothrix ku80* |
| **35** | GGAGCTCGACAACAAGGGTC | *P. brasiliensis* | Deletion control of *Sporothrix ku80* |

### Supplementary Figure 1

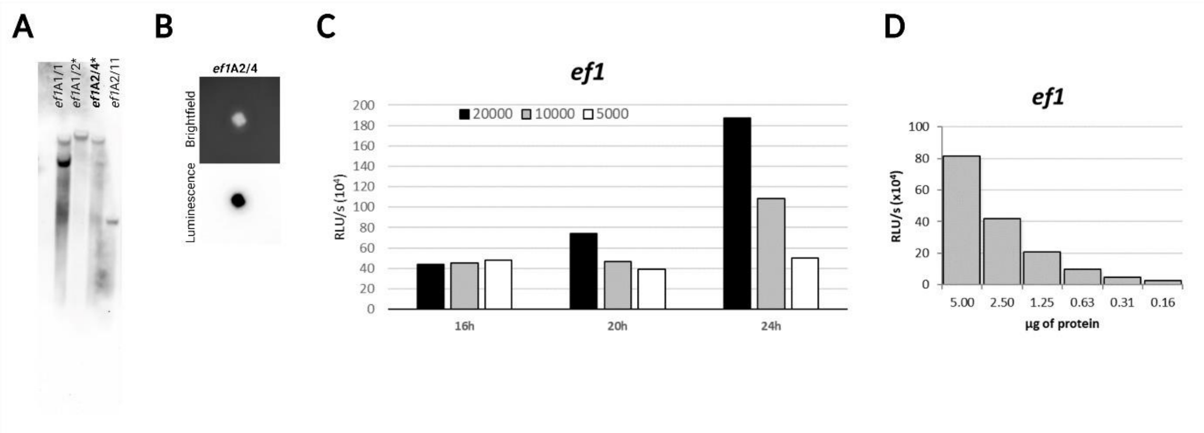


**Supplementary Figure 1:** Luciferase construct for expression in *Sporothrix* species. The efficiency of the plasmid harbouring *PbLuc* under the control of elongation factor 1-gamma (*ef1*) promoter from *Paracoccidioides* was tested in *A. niger* transformation*.* **(A)** *A. niger* transformant containing single integration of the plasmid was selected by Sothern blot analysis. **(B)** 500 spores from *A. niger PbLuc* transformant containing single integration of the plasmid were spotted in GG10 containing D-luciferin (0.2 mM). Brightfield (upper panel) and luminescence (bottom panel) were recorded after 2days of incubation at 28 $^{o}$C. **(C)** *In vivo* and **(D)** *in vitro A. niger* assay for *PbLuc* expression under the control of *ef1* promoter.

**Supplementary Figure 2**

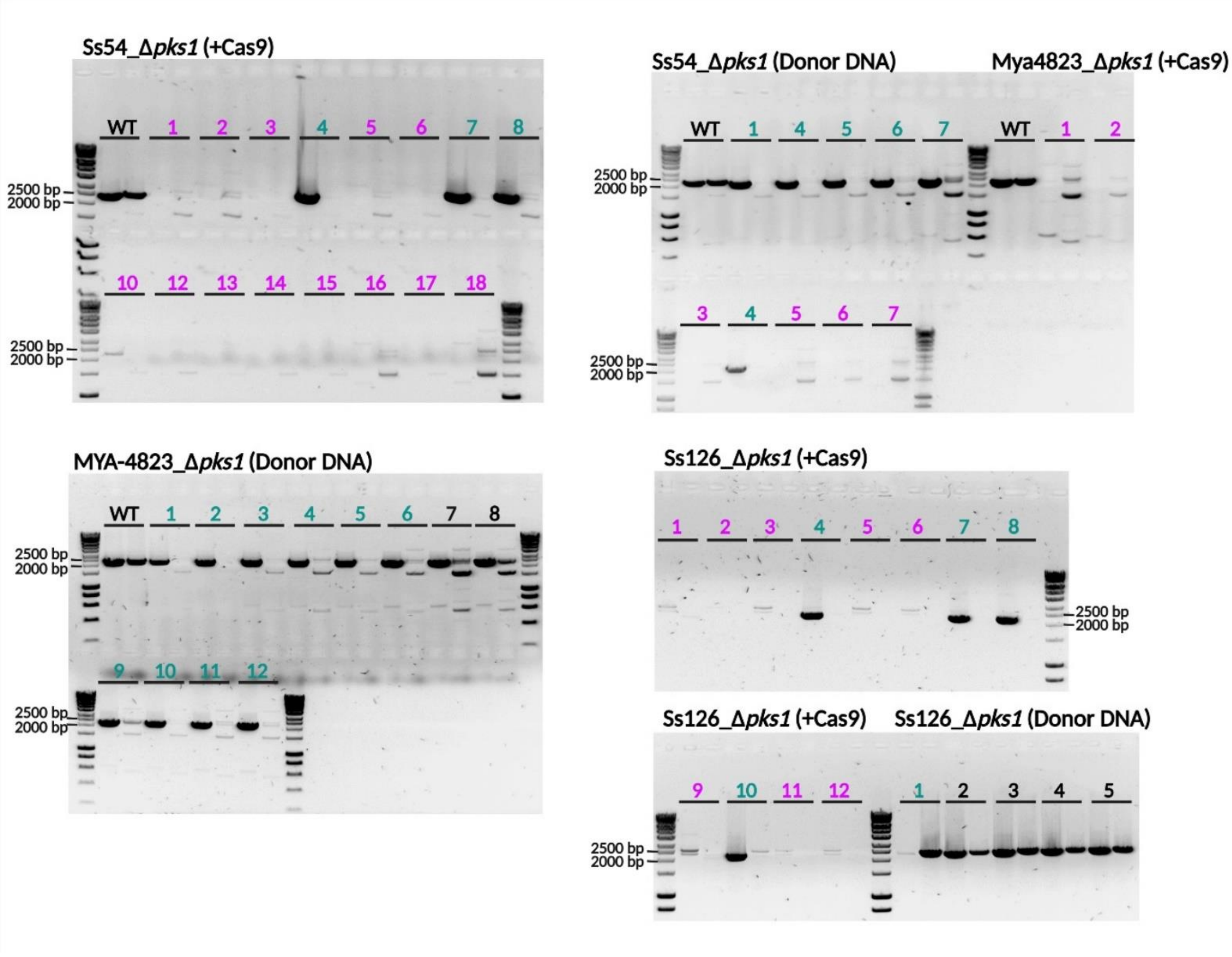


**Supplementary Figure 2:** Validation of *pks1* disruption. PCR analysis for identifying the disruption of *pks1* coding region using the gDNA from 16 Ss54 colonies, 7 MYA-4823 colonies and 12 and Ss126 colonies. For each transformant the left-hand lane represents PCR diagnosis of the *pks1* deletion at the upstream gRNA site, and the right-hand lane the diagnosis of the *pks1* deletion at the downstream gRNA site. Strains with the WT coding region (black) yielding PCR products at 2257 bp and 2337 bp or containing one PCR product (green) were excluded from the subsequent analysis. Samples double negative (purple) for the presence of *pks1* coding region were further analysed for the integration of deletion cassette (Figure 2C).

**Supplementary Figure 3**

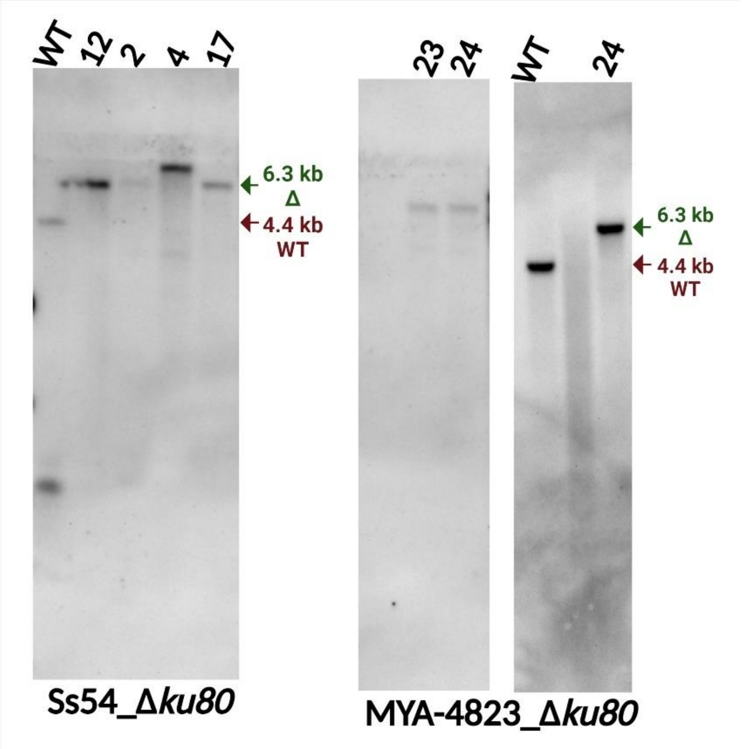


**Supplementary Figure 3:** Southern blot analysis of *ku80* deletion strains. Southern blotting showing 4 and 2 transformants from Ss54 and MYA-4823 containing in locus integration. Diagnostic bands of 4.4 kb for wild type (WT) and 6.3 kb for a deletion mutant (Δ) are highlighted.

**Supplementary Figure 4**

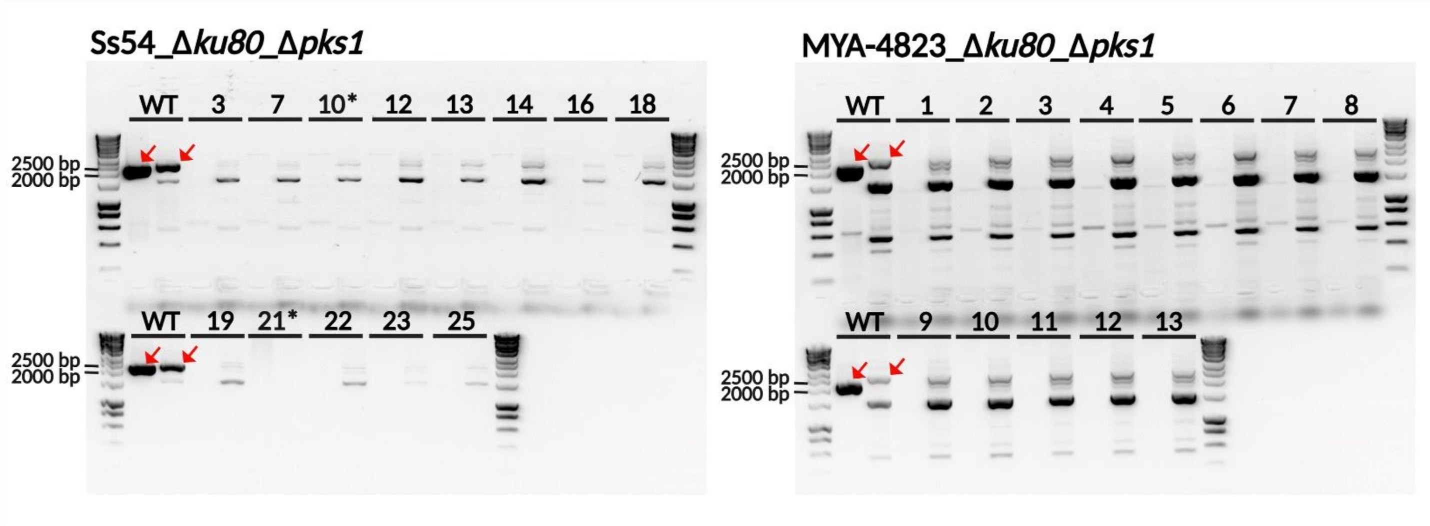


**Supplementary Figure 4:** Validation of *pks1* disruption of the double deletion transformants. PCR analysis for the detection of coding region using the gDNA from 13 MYA-4823_Δ*ku80*_Δ*pks1* and Ss54_Δ*ku80*_Δ*pks1* transformants. Strains with the WT coding region yield PCR products at 2257 bp and 2337 bp (red arrows). All transformants did not present specific PCR products of the expected size and were analysed for the integration of nourseothricin resistance cassette (Figure 4A).

### Supplementary Methods

***Aspergillus* transformation and luciferase assay.** *Aspergillus niger* A1144 Δ*pyrG* (Geib et al., 2019) was cultivated at 28 °C in modified *Aspergillus* Minimal Medium-GG10 (Geib et al., 2016) supplemented with 10 mM uridine and 2% (w/v) agar when required. For DNA extraction and luciferase activity assay, *A. niger* was cultured overnight YPD medium. A1144 Δ*pyrG* conidia were harvested from slants in Phosphate-Buffered Saline (PBS) with 0.1% (v/v) Tween 20, washed twice with PBS and the spore concentration determined by counting in a haemocytometer chamber. The protoplast-mediated transformation was performed as previously described using p*ef1*:*Pbluc*$_{OPT\_red}$:t*eno1*_URABlaster_pUC19 (Geib *et al.*, 2019; Geib *et al.*, 2016; Milhomem Cruz-Leite et al., 2022). To identify transformants containing single integration of the plasmid, we performed Southern blot analyses of the *Eco*RI digested gDNA for the detection of *PbLuc* as previously described (Milhomem Cruz-Leite *et al.*, 2022). 500 conidia of the transformant *ef1*A2/4 from *Aspergillus niger* was spotted in GG10 agar supplemented with 0.4 mM of D-luciferin and incubated at 28 °C for 2 days. The bioluminescence was recorded using a ChemiDoc XRS+ system (Bio-Rad). The *PbLuc* activity was measured in crude cell-free extracts and *in vivo* as described by (Milhomem Cruz-Leite *et al.*, 2022).
